# High-Resolution, Multidimensional Phylogenetic Metrics Identify Class I Aminoacyl-tRNA Synthetase Evolutionary Mosaicity and Inter-modular ‘Coupling

**DOI:** 10.1101/2020.04.09.033712

**Authors:** Charles W. Carter, Alex Popinga, Remco Bouckaert, Peter R. Wills

## Abstract

The provenance of the aminoacyl-tRNA synthetases (aaRS) poses unusually challenging questions because of their role in the emergence and evolution of genetic coding. We investigate evidence about their ancestry from highly curated structure-based multiple sequence alignments of a small “scaffold” that is structurally invariant in all 10 canonical Class I aaRS. Statistically different values of two uncorrelated phylogenetic metrics—residue by residue conservation derived from Clustal and row-by-row cladistic congruence derived from BEAST2—suggest that the Class I scaffold is a mosaic assembled from distinct, successive genetic sources. These data are especially significant in light of: (i) experimental fragmentations of the Class I scaffold into three partitions that retain catalytic activities in proportion to their length; and (ii) multiple sources of evidence that two of these partitions arose from an ancestral Class I aaRS gene encoding a Class II ancestor in frame on the opposite strand. Two additional metrics output by BEAST2 vary in accordance with the presumed functionality endowed by the various modules. The new evidence supplements previous aaRS phylogenies. It identifies a previously characterized 46-residue Class I “protozyme” as preceding the adaptive radiation of the superfamily containing variations of the Rossmann dinucleotide binding fold related to amino acid discrimination, and thus as root of that molecular tree. Such a rooting is consistent with near simultaneous emergence of genetic coding and the origin of the proteome, resolving a conundrum posed by previous inferences that Class I aaRS evolved long after the genetic code had been implemented in an RNA world. Further, it establishes a timeline for the growth of coding from a binary amino acid alphabet by pinpointing discontinuous enhancements of aaRS fidelity.

**Author Summary:** Phylogenetic analysis uncovers evolutionary connections between different protein superfamily members. We describe complementary, uncorrelated, phylogenetic metrics that support multiple evolutionary histories for different segments within members of the Class I aminoacyl-tRNA synthetase superfamily. Using a carefully curated 3D crystal structure superposition as the primary source of the multiple sequence alignment substantially reduced dependence of these metrics on empirical amino acid substitution matrices. Two metrics are derived from the amino acid distribution observed in each successive position. A third depends on how individual sequences distribute into phylogenetic tree branches for each of the ten amino acids activated by the superfamily. All metrics confirm that a segment previously identified as an inserted element is, indeed, a more recent acquisition, despite its structural conservation. The residue-by-residue conservation metrics reveal significant co-variation of mutational frequencies between a core segment that forms the amino acid binding site and a neighboring segment derived from the more recent insertion element. We attribute that covariation to the differentiation of superfamily members as evolutionary divergence enhanced amino acid specificity. Finally, evidence that the insertion element is a recent acquisition implies a new branching order for much of the proteome.

## Introduction

The emergence of the aminoacyl-tRNA synthetases, aaRS, is a quintessential chicken and egg puzzle whose solution would demystify the origins of coded protein synthesis. How did aaRS enzymes gain the reflexive property of collectively being able to use relationships in the universal genetic code to convert the sequences of base triplets in their own genes into functional amino acid sequences that make the code work? The detailed trajectory by which genes for the two essential superfamilies appeared, grew to acquire catalytic proficiencies and radiated to refine their dual amino acid and cognate tRNA specificities is thus a crucial chapter in the book of life.

Both aaRS Classes have separate catalytic and anticodon-binding domains. Only the catalytic domains within each superfamily share the same architectures [1]. In this paper, we develop and analyze high-resolution phylogenetic metrics relevant to the mosaic evolutionary histories of those shared architectures. Anticodon-binding domains are idiosyncratic and not considered here.

Phylogenetic analyses of the Class I aaRS superfamily [2-8] generally agree that clades for each amino acid are monophyletic, divide into three subclasses [1, 9], including IleRS, ValRS, LeuRS, and MetRS (Subclass IA); GluRS and GlnRS (Subclass IB); and TyrRS and TrpRS (Subclass IC). Restricting analysis to the bacterial aaRS as they point more faithfully to the root of protein synthesis [10], we exclude the archaeal Subclass IB Class I LysRS. CysRS and ArgRS families, assigned originally to Subclass IA [9], are difficult to assign, with one or the other appearing instead with Subclass IB [for example, see 5]. A parallel sub-classification in Class II aaRS strengthens connections [11-13] between Class I and II phylogenies and the physical chemistry differentiating the 10 amino acids in each Class.

### Three segments within the Class I aaRS catalytic domains suggest a succession of intermediate states dating from well before the Last Universal Common Ancestor

Genetic deconstruction of Class I aaRS modules [14] identified a core whose ∼130 residues form a nearly intact active site, and which experiments show retain a major fraction of full-length aaRS catalytic proficiency—estimated as the transition-state stabilization free energy—for both amino acid and tRNA substrates [15-17] (Fig 1). Because of their high catalytic proficiency in both essential reactions, these and similar cores derived from Class II aaRS were called “Urzymes”, from the German prefix “Ur” = original, authentic. Class I and II Urzymes differentiate between amino acids from the two sets of substrates, with Class I Urzymes activating Class I, in preference to Class II amino acids by ∼1.0 kcal/mole, and conversely [18-20].

**Fig 1.**
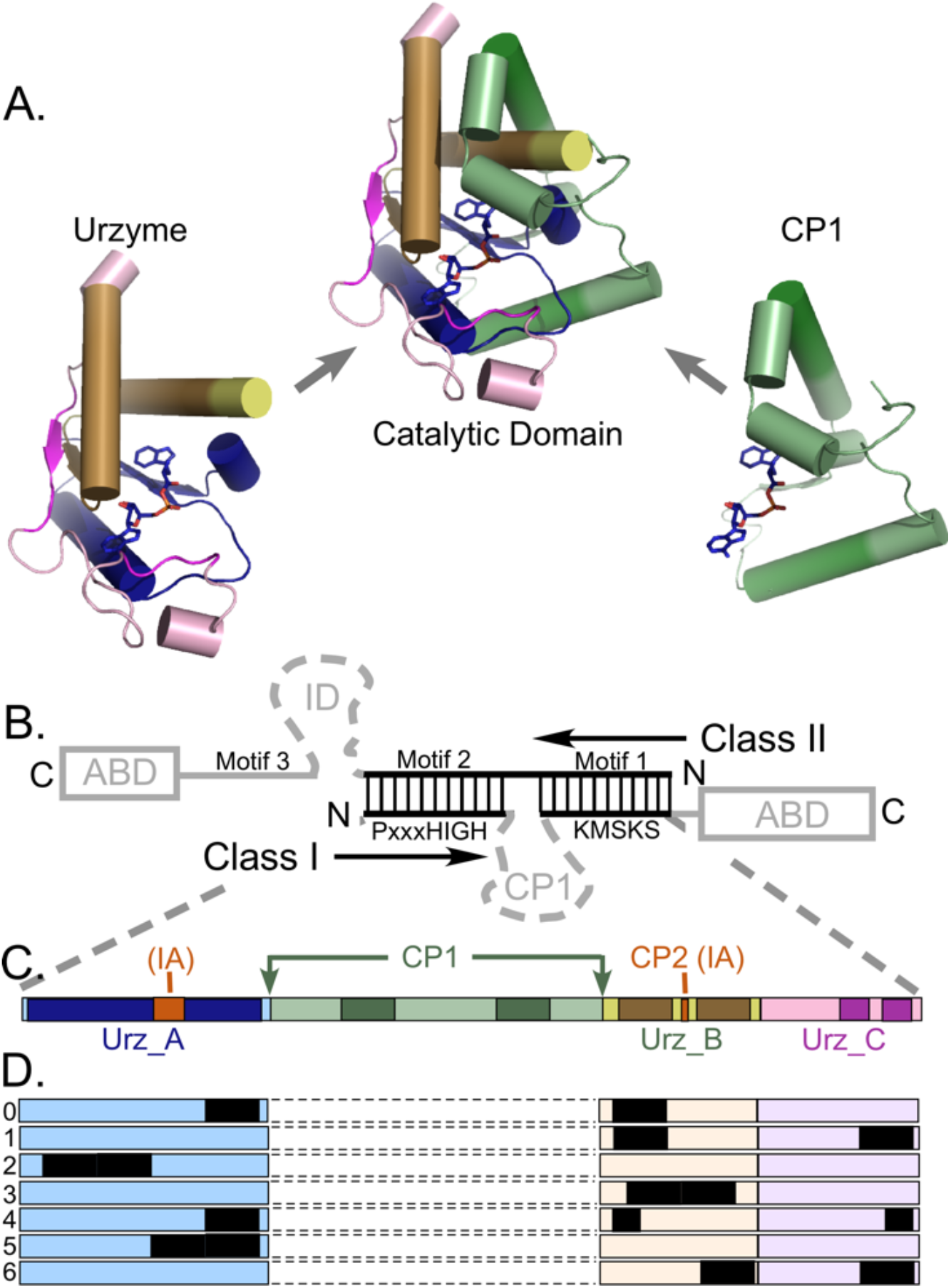
Structural modules underlying the hypothesis and organization of data adduced as evidence. **A**. The Class I Urzyme and internal CP1 insertion that together make up the Class I catalytic domain. Cartoons were prepared with Pymol [75] from coordinates (PDB ID 1I6L) for the tryptophanyl-tRNA synthetase, the smallest Class I aaRS. The activated aminoacyladenylate is drawn as sticks. The primary size variation the remaining nine Class I aaRS occurs as further insertions within CP1 as structures displayed here are essentially unchanged. Anticodon-binding domains are idiosyncratic, and not shown. **B**. Overlapping portions of Class I and II aaRS as envisioned in an ancestral bidirectional gene [33] coincide with the respective Urzymes [19, 30]. Vertical lines denote ancestral base-pairing. Grey segments were presumably more recent additions. In particular, insertion of CP1 is incompatible with bidirectional coding. **C**. Schematic location of CP1 between two roughly equal fragments of the Urzyme, colored to allow identification of structural fragments in **A**. Intensely colored segments are highly conserved secondary structures in all 10 Class I aaRS and compose the Class I scaffold. Subclass I A enzymes contain one or more additional insertions (red). **D**. Sampled alignments consisting of different segments totaling 20 amino acids, selected as described in Methods. **C** and **D** amplify the Class I base-paired portion of **B**. The protozyme (Urz_A) is the amino terminal β-α-β crossover connection (blue), the amino acid binding pocket (Urz_B) is formed by the protozyme and two intermediate α-helices (amber). The KMSKS loop (Urz_C) is rose. Previous work [25] established that the CP1 insertion (green) has little impact on the enzymatic properties of the TrpRS Urzyme unless the anticodon-binding domain is also present. Colors match those in (**A**).

Even smaller 46 residue subsets from both Class I and II retain nearly half the full catalytic proficiency in the critical amino-acid activation reaction [21], whose uncatalyzed rate is rate-limiting for protein synthesis. These primitive modules were called “protozymes”, from the Greek πρoτo = first, to distinguish them from Urzymes. The Class I protozyme coincides with the N-terminal crossover connection of the Rossmann fold, which contains a distinctive 3D packing motif that recurs in ∼25% of the proteome [22] and imposes distinct, functionally relevant conformational states tightly coupled to catalysis in TrpRS [23-29].

Fully active Class I and II protozymes have been expressed from a single gene designed to encode their structures on opposite strands. Experimental Michaelis-Menten parameters of all four protozymes—Class I and II; native and bidirectional—are, remarkably, the same within experimental error [21]. Experimental [21, 30, 31] and bioinformatic evidence [10] therefore support the hypothesis of Rodin and Ohno [32, 33] that the two aaRS Classes descended from opposite strands of a single bidirectional gene.

The progressive biochemical functionality of aaRS protozymes, Urzymes, and catalytic domains; the fact that their structures are universally conserved within both Classes; and the evidence that they descended from one bidirectional ancestral gene; all imply that they are legitimate experimental models for ancestral evolutionary aaRS forms that participated throughout the emergence of genetic coding and well before LUCA [14]. Moreover, to date, no such sequential intermediates have been derived for other superfamilies. Thus, more appears to be known about the modular evolution of the two aaRS superfamilies than for any other ancient superfamily, making the Class I aaRS a highly relevant subject for this work.

### A variable-length insertion interrupts Class I aminoacyl-tRNA synthetase catalytic cores

Class I aaRS catalytic domains contain a variable-length insertion called “connecting peptide 1” or CP1 [34, 35], which is detailed in Fig 1. CP1 insertions include the large editing domains of the aaRS for aliphatic amino acids, and thus account for the variable size of Class I catalytic domains. CP1 always intersects the Rossmann fold immediately after the Protozyme between structurally homologous residues that are ∼4.5 Å apart and can be replaced by a peptide bond without structural disruption to produce the Class I Urzyme [30].

### The scaffold identifies invariant secondary structures outside the Urzyme

Evidence that the essential catalytic architectures observed in aaRS Urzymes arose from a bidirectional gene implies, *ipso facto*, that sequences inconsistent with bidirectional coding represent accretion of new genetic material. In this paper we address an apparent contradiction between the Rodin-Ohno hypothesis and extended conservation into the CP1 segment. Independent 3D superposition of aaRS crystal structures [2, 30, 36] revealed considerable structural conservation within the catalytic domains of all ten members of each Class—see Fig 1 of [36]. Structural homology across the Class I superfamily extends beyond the Urzyme, into CP1. We call the secondary structures within these conserved cores “scaffolds”.

The CP1 insertions are incompatible with, and their introduction would necessarily have ended, bidirectional genetic coding of ancestral Class I and II aaRS (Fig 1B). Class I Urzyme boundaries delimited decisively by potential bidirectional coding of Class II Urzymes constitute only about 80% of the Class I scaffold; the remainder consists of 10-residue α-helical segments near the beginning and end of the CP1 insertion (Fig 1). Thus, if the Rodin-Ohno hypothesis is correct, then the CP1 insert must derive from a distinct genetic source.

Data summarized in the three previous sections lead us to expect metrics derived from MSAs for modules with distinct ancestries to differ quantitatively. For these reasons, we developed new, accurate ways to quantify the phylogenetic signatures of the protozyme, Urzyme, and CP1 aaRS segments in order to assess the likelihood that CP1 represents a more recent acquisition by ancestral Urzymes.

### Novel applications of phylogenetics

This work introduces four novelties that, together, reinforce the conclusion that modular components of the ancestral aaRS have different genetic histories:

i. We threaded sequences into closely-related crystal structures to assemble multiple sequence alignments (MSAs) based on three-dimensional structure superposition, to avoid depending on amino acid substitution matrices to define equivalent positions.
ii. We develop multi-dimensional phylogenetic metrics from the ensemble of phylogenetic trees obtained by Markov Chain Monte Carlo (MCMC) simulations to compare segments of the MSA. We show that these three metrics are uncorrelated, and demonstrate their statistical significance by multiple regression.
iii. We increase the effective analytical resolution by applying the metrics to partitions of the MSA that have been extensively characterized experimentally, providing novel insight into the functional modularity of the superfamily and molecular phylogenetic support that they are successive evolutionary intermediates in the generation of the Class I aminoacyl-tRNA synthetase superfamily.
iv. We identify key two-way interactions between mutation rates in different MSA segments. Both involve the amino acid binding site and one is central to the enhancement of amino acid specificity enabled by the CP1 insertion.

These results suggest important modifications of conventional evolutionary scenarios for major parts of the proteome containing Rossmann dinucleotide fold domains, and strengthen the proposal that the genetic code development is coupled intimately to the structural evolution of aaRS.

## Results

Phylogenetic metrics derived from trees constructed for different subset MSAs—CP1; the Urzyme; and its three distinct modules Urz_A (the protozyme), Urz_B (the amino acid binding site, and Urz_C (the PP_i_ binding site)—furnish unprecedented insight into the genetic modularity of the Class I aaRS. In summary, they suggest that CP1 is derived from a substantially more recent and less cladistically coherent genetic entity, as previously posited [30]. Similarly, they suggest that the protozyme may be from an older genetic entity.

### CP1 has lower and more uniform apparent site-to-site mutation rates between aaRS for different amino acids

The BEAST 2 MCMC algorithm tracks the extent of site-to-site evolutionary of sequence variation in using the Tree height and Shape metrics described in Methods. CP1 sequences from the scaffold exhibit much reduced mutation rates and variance, consistent with their being a more recent evolutionary acquisition. Sequences within the Class I Urzymes exhibit higher and more variable mutation rates, consistent with greater adaptive radiation, hence deeper ancestry than CP1 sequences.

The Tree height metric is summarized in (Fig 2A, B). Sequences responsible for the elevated Urzyme Tree height are identified by regression against the independent parameters of the design matrix (Table 1) in Fig 2C. Tree height = 2.17 - 1.4 * *Segment B* + 3.2 * *Urzyme* - 2.7 * *protozyme* + 1.7 * *Segment B* * *protozyme*. All β coefficients are significant with P-values < 0.005. Foremost among the positive contributors is the Urzyme. However, the interaction between the Protozyme (segment A) and segment B is also quite significant. We note that CP1 sequences exhibit substantially smaller Tree height and variance (i.e. higher Shape), consistent with it being a more recent genetic entity.

**Table 1.** Design matrix for regression analysis^1^. ^1^ Shaded columns are dependent variables; unshaded columns are independent variables and serve as predictors. Numerals in columns A, B, C, and CP1 are proportional to the total number of amino acids in the respective MSAs. Two additional independent variables, Urzyme and protozyme, constructed in a related fashion, are not shown. NUMB is the number of amino acids in the alignment.

| Subset | <S><sub>WAG</sub> | <S><sub>LG</sub> | <Q> | Tree height | Shape | A | B | C | CP1 | NUMB |
| --- | --- | --- | --- | --- | --- | --- | --- | --- | --- | --- |
| Full | 0.9 | 0.86 | 48.8 | 3.69 | 1.17 | 4 | 3 | 1 | 2 | 103 |
| Urzyme | 0.87 | 0.82 | 52.4 | 3.40 | 1.60 | 4 | 3 | 1 | 0 | 83 |
| Protoz | 0.85 | 0.85 | 52.3 | 2.71 | 1.38 | 4 | 0 | 0 | 0 | 46 |
| CP1 | 0.34 | 0.35 | 33.7 | 2.17 | 4.49 | 0 | 0 | 0 | 2 | 20 |
| Urz_0 | 0.76 | 0.73 | 50.1 | 3.03 | 1.46 | 1 | 1 | 0 | 0 | 20 |
| Urz_1 | 0.71 | 0.68 | 57.8 | 3.99 | 0.91 | 0 | 1 | 0.5 | 0 | 20 |
| Urz_2 | 0.72 | 0.7 | 55 | 2.78 | 1.13 | 2 | 0 | 0 | 0 | 20 |
| Urz_3 | 0.62 | 0.62 | 45.6 | 2.54 | 2.26 | 0 | 2 | 0 | 0 | 20 |
| Urz_4 | 0.82 | 0.77 | 58.5 | 2.41 | 1.21 | 2 | 1 | 0.5 | 0 | 20 |
| Urz_5 | 0.78 | 0.8 | 45.6 | 2.88 | 1.98 | 4 | 0 | 0 | 0 | 20 |
| Urz_6 | 0.62 |  | 56.6 | 2.65 | 1.13 | 0 | 2 | 1 | 0 | 20 |

**Fig 2.**
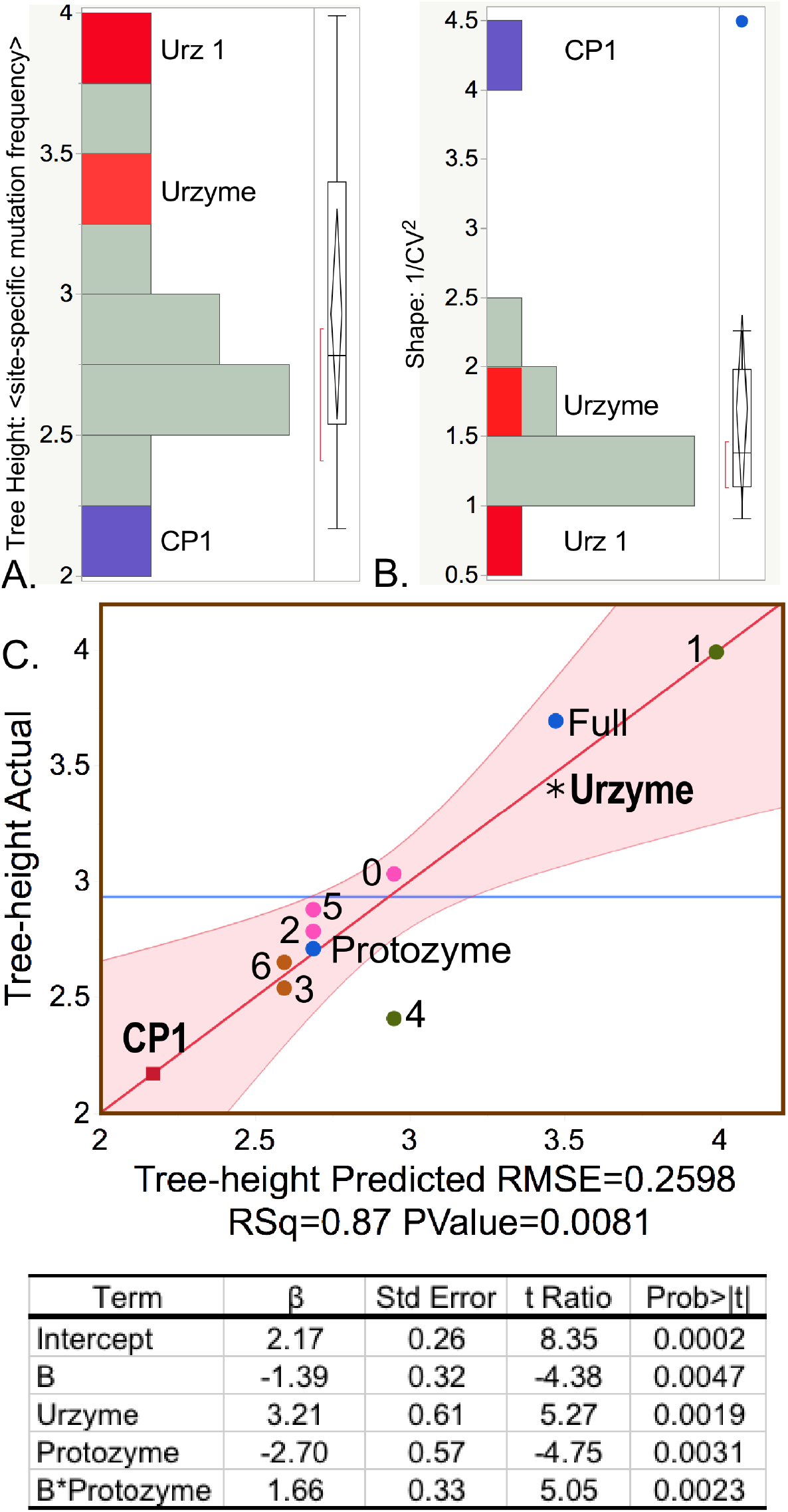
Histograms of the Tree height (**A**) and Shape (**B**) metrics highlight differences between sequence variability within CP1 (blue) and Urzyme (red) sequences. The Tree height is the reciprocal of the estimated mutation rate per site. CP1 has the lowest site-specific mutation rate and the highest Shape parameter (blue dot implies statistical significance). The Urz_1 subset of amino acids includes a stretch of 10 amino acids along the specificity determining helix, where the highest site-specific mutation frequency occurs within the Urzyme. **C**. Regression model showing the dependence of Tree height on MSA subsets. The horizontal blue line in this and other regression curves is the average value of the dependent variable. Dots represent the various MSAs in the design matrix and are colored and distinguished by different symbols for identification. R^2^ for this model is 0.87. Numerals refer to the 20 residue subsets defined in Fig 1. The F-ratio is 9.9 with a P-value of 0.008.

The negative logarithm of the Shape parameter is highly correlated with the conservation quality, ⟨*Q*⟩, defined by Clustal [37, 38] (Fig 3A), lending intuitive insight into the physical meaning of the latter. The designation “Conservation Quality” is apparently misleading in suggesting that the significantly smaller ⟨*Q*⟩ value for the CP1 MSA (Fig 3B) implies that it is less well conserved, in apparent conflict with its reduced Tree height. In fact, the colinearity of –log_10_(Shape) and ⟨*Q*⟩ led us to pursue the equivalence between the two metrics. Fig 3C, D show that the two metrics depend in nearly identical fashion on predictor columns from the design matrix in Table 1. It appears then that the ⟨*Q*⟩ metric does not measure mutational variation itself, but rather the inverse of its variance.

**Fig 3.**
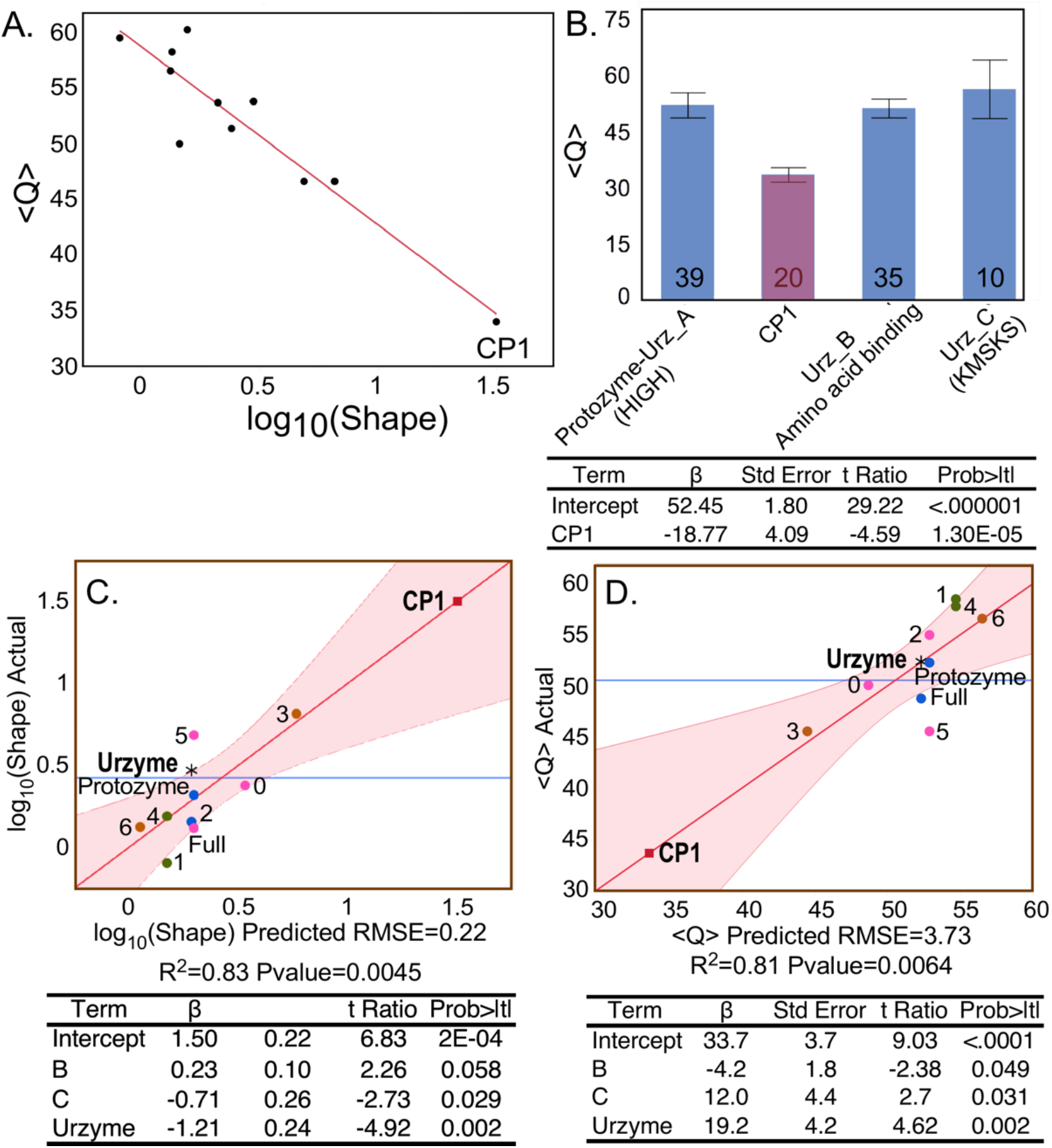
Position-specific metrics. **A**. Colinearity of the logarithm of the Shape parameter of the gamma distribution of the Tree heights estimated by BEAST2 for different MSAs derived from the Class I scaffold ⟨*Q*⟩, and the mean conservation quality scores, ⟨*Q*⟩ [38] obtained directly from respective MSAs. **B**. The conservation quality, <Q>, for the four segments of the Class I scaffold alignment. Class I signature peptides [1] are in parentheses. Error bars show the standard error of the mean over all positions within the segment. The numbers of amino acids in each segment are given for each histogram. **C**. and **D**. are regression models showing nearly identical dependence of log_10_(Shape) and ⟨*Q*⟩, respectively, on predictors in the design matrix (Table 1). Dots represent different MSAs. They are labeled and colored as in Fig 2 to emphasize the extraordinary similarity of the two site-by-site metrics derived two different ways.

Moreover, the extended similarity between Fig 3C and Fig 3D suggest that the site-by-site metrics (Tree height and Shape) can provide quite high resolution evidence on the evolution of modularity. Regression of Shape on the independent parameters of the design matrix in Table 1 (Fig 4) resulted in the unique model: Shape = 1.37 + 0.4 * *Segment B* − 1.08 * *Segment C* + 1.56 * *CP*1 − 0.57 * *Segment B* * *CP*1. All β coefficients are significant and two have P-values < 0.005.

**Fig 4.**
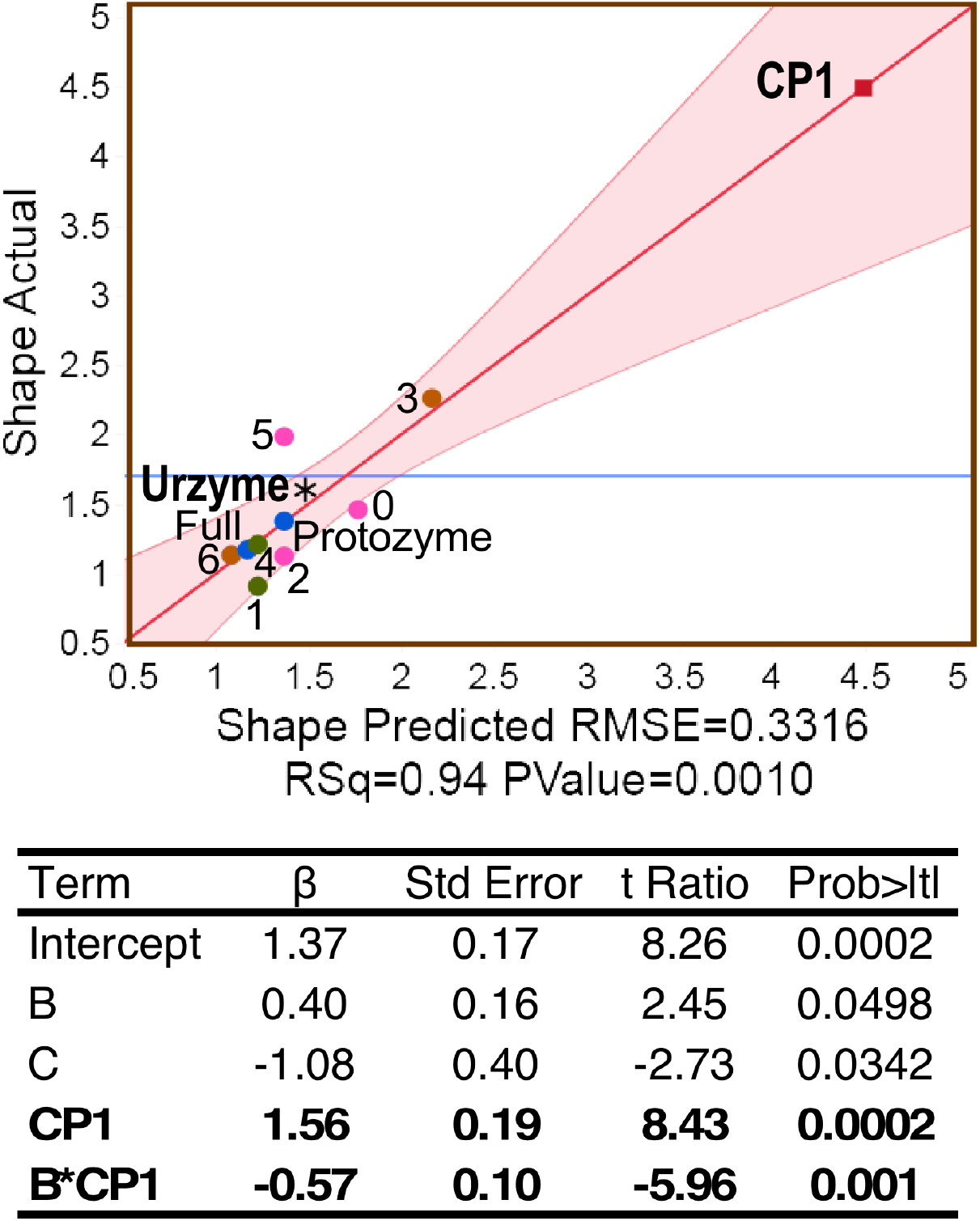
Shape parameter dependence. **A**. Plot of the multivariate regression model for Shape. R^2^ = 0.94 for the regression and the P value for the F ratio is 0.0006. **B**. Regression coefficients and their Student t-test probabilities. Points for trees constructed using both WAG and LG substitution matrices are shown for comparison. Significant predictors of Shape are the amino acid positions in CP1, those in the B-fragment containing the amino acid specificity-determining helix (sand; Fig 1A), and their two-way interaction.

### Class I Urzyme- and CP1-based trees form distinctly different clades

Urzyme and CP1 partitions of the MSAs produce substantially different trees (Fig 5A). In particular, although all ten Class I aaRS clades are monophyletic in the trees for the Urzyme MSAs, three enzymatic clades in the CP1 trees—MetRS, ValRS, and TyrRS—are polyphyletic. Moreover, the Urzyme clades are constrained by tight envelopes, whereas the CP1 envelopes are poorly defined.

**Fig 5.**
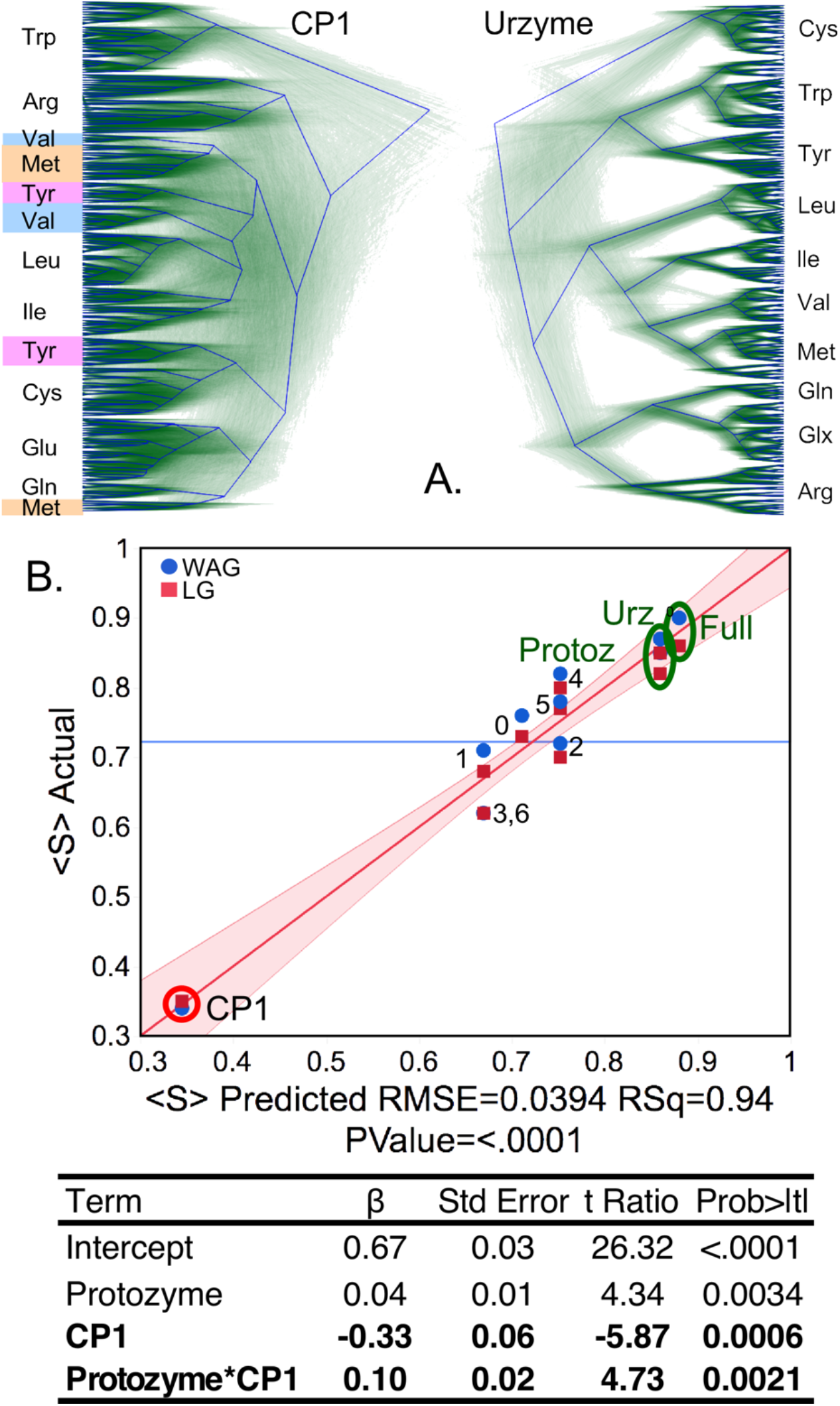
Phylogenetic support. **A**. DensiTree representations of the Urzyme and CP1 segments of the Class I structural scaffold. Each aaRS is monophyletic in the Urzyme alignment, whereas several of the clades in the CP1 alignment, highlighted in color, have multiple ancestries. **B**. Regression model for ⟨*S*⟩ _i_. Dots are colored and labeled as in Fig 3. Values in the Regression table are only for the WAG matrix, as use of both WAG and LG matrices unduly enhances the P-values of the β-coefficients. The R^2^ and P value of the model’s F-ratio are shown under the X-axis. Coefficients and their statistics for the model are given in Table 2. Individual ⟨*S*⟩ values are labeled to enable comparison with Fig 1C. R^2^ was 0.94 for 20 observations, and the F-ratio for the regression table was 92.4 with a P-value <0.0001.

Support, ⟨*S*⟩_*i*_, the fraction of all trees for which each aaRS clade, *i*, is monophyletic, was averaged over all Class I aaRS types to give the mean support, ⟨*S*⟩ = Σ*S_i_*/10. That operation was repeated for trees built for the full scaffold MSA, (Full), the Urzyme, CP1, protozyme (segment A in Fig 1), and seven subset MSAs each consisting of twenty amino acids in blocks of five residues as described in Methods. For the eleven rows of the design matrix (Table 2), we computed ⟨*S*⟩ for populations of trees constructed using the conventional WAG [39] substitution matrix and repeated using the more recent LG [40] substitution matrix used in [8].

**Table 2.**
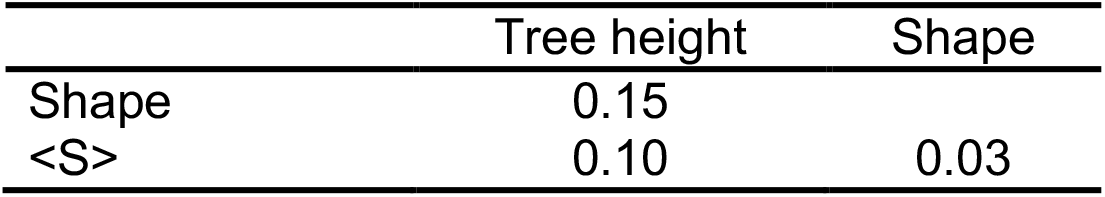
Cross-correlation R^2^ values between phylogenetic metrics.

|  | Tree height | Shape |
| --- | --- | --- |
| Shape | 0.15 |  |
| <S> | 0.10 | 0.03 |

Contributors to the variance of ⟨*S*⟩ were assessed using stepwise multiple regression, which resulted in the unique model: ⟨*S*⟩ = 0.67 + 0.04 * *protozyme* − 0.33 * *CP*1 + 0.1 * *protozyme* * *CP*1 summarized in Fig 5. All three coefficients are significant at better than the 99% confidence level, with P-values < 0.005. As suggested by its position in the regression curve in Fig 5, trees built from CP1 residues have significantly less support than those from the Urzyme or any of its 20-residue subsets. The A segment coincides with the protozyme MSA; its dominant impact on the variance of ⟨*S*⟩, contributing positively both via its intrinsic effect and by its two-way interaction with CP1, is consistent with its being close to the root of the Class I aaRS superfamily tree.

Thus, although the 3D structures from the Urzyme and CP1 partitions have comparable structural homology, they have markedly different phylogenetic signatures.

### Phylogenetic metrics—Tree height, Shape, and Support—reveal significant, high-resolution genetic mosaicity

Our quantitative evidence probes far deeper into evolutionary time than previous phylogenetic analyses of Class I aaRS [3, 7, 41-43]. That depth both calls for caution and is a source of great interest. Constructing aaRS trees is fundamentally ambiguous because at each node in any conceivable tree, the coding alphabet and dimension of the operational substitution matrix necessarily both change by integer steps as Class I and II trees branch into multiple families. Trees for the two superfamilies are thus necessarily interdependent, so that the dimensions of all possible substitution matrices start from 2 and end at 20. Thus, it is uncertain what should be inferred from phylogenetic metrics for a single superfamily.

These difficulties are substantially offset by the consistency of site-by-site and row-by-row metrics with the construction and experimental characterization of aaRS protozymes [21] and Urzymes [14, 15, 18, 19, 24, 25, 31] (Fig 1). Such deep evolutionary intermediates are, at present, manifestly unique to the aaRS. The extensive experimental and structural context of that consistency strengthens our conclusions even without comparable analysis of Class II aaRS, now in progress, especially in light of the following observations.

#### The three metrics are uncorrelated

The CP1 MSA is a substantial outlier for all three metrics, suggesting that the metrics may be correlated. Removing the CP1 entries from the design matrix eliminates any correlation between Tree height, Shape, or ⟨*S*⟩, Table 2. The three types of metrics are therefore essentially uncorrelated and provide independent insights.

#### Differences between Urzyme and CP1 sequences are statistically meaningful

If the Class I aaRS sequence partitions compared in Table 1 were all drawn from continuously replicated ancient genetic sources, subject to comparable selection history since their emergence, the null hypotheses would produce similarly conserved sequences and comparably congruent trees for the Urzyme and CP1 partitions. The log-worth values (i.e., –log(P)) for the higher mutational frequency (Tree height; 2.7), its variance (Shape; 3.3) and lack of congruence for phylogenetic trees, (⟨*S*⟩ ; 3.4), imply with high statistical significance that all metrics for CP1 and Urzyme sequences are drawn from different populations, corroborating the argument—based on bidirectional coding ancestry— that the CP1 sequences represent genetic information acquired more recently by the Urzymes.

#### The numbers of residues in different segments drawn from the MSA have no detectable impact on any phylogenetic metric

One might suppose that degraded congruence of the CP1 trees results from the fact its MSA has only ∼0.25 as many amino acids as the Urzyme MSA. The insignificant impact of NUMB on the regression models for any score and the clustering of the 20-residue ⟨*S*⟩ values with the intact Urzyme MSA (Fig 5) confute that expectation.

#### Threading does not force any particular comparison between different aaRS types

Threading increased the reliability of structure-based alignments within any aaRS type by adding sequences. Scaffold positions were rigorously defined as structural homologs from the close proximity of their alpha carbon coordinates in multiple PDB structure alignments (i) among aaRS for any single amino acid, drawn from very diverse bacterial species, that only then (ii) produced a grand scaffold MSA across all Class I aaRS types. Additions produced by threading thus have only second-order effects on structural superpositions of aaRS for different amino acids, adding precision to our analysis without overtly influencing choices on which our conclusions depend.

#### Distinguishing features of CP1 sequences are evident without considering indels

Figs 2-5 together illustrate a comprehensive, near-optimally quantitative three-dimensional comparison of partitions in the Full Class I aaRS scaffold MSA. The crux of what the data suggest is that CP1 is more recent than the Urzyme, its evolutionary divergence and variance—evidenced by its Tree height and Shape—is much reduced, but its phylogenetic consistency—evidenced by ⟨*S*⟩ —is much reduced, relative to the corresponding metrics for Urzyme segments. This conclusion is evident, from sequences with strict 1-1 correspondences between 3D crystal structures, excluding the large and variable-length indels that dominate CP1 insertions in most aaRS types.

#### Neither convergent evolution nor horizontal gene transfer are likely explanations for the Urzyme/CP1 distinction

The congruent clade structure of Urzyme-derived sequences from the scaffold fall into consensus groupings characterizing Class I aaRS as a coherent superfamily and does not require mechanisms beyond mutation and selection from a single common ancestor. The aberrant behavior of sequence variations in the CP1 insertion (Figs. 2-5) might suggest appeal to such processes. Treatments of Class I aaRS evolution based on full MSAs [6, 44, 45] show evidence of horizontal gene transfer (HGT), genetic transpositions and large scale insertion/deletion events, of which CP1 is the foremost example.

Wherever full-length bacterial Class I enzymes with a particular amino acid substrate specificity are represented by more than one canonical structure, as has been described for IleRS and MetRS [see Fig. 3 in 44], that bipartite distribution in genome space is adequately explained in terms of early HGT into bacteria from an archaean/eukaryotic ancestor, but not in terms of convergent evolution. CP1 sequence disparities at homologous sequence positions in TyrRS, MetRS, and ValRS behave in the opposite manner: the higher variance of their ⟨*Q*⟩ values producing lower ⟨*S*⟩ values by allowing their evolutionary paths to wander widely, often crossing in sequence space, instead of forming multiple well-defined clades reasonably distant from one another in sequence space that would suggest HGT.

### Phylogenetic metrics have functionally relevant interpretations

Regressions in Fig 3 were performed to demonstrate equivalence of the –log10(Shape) and the Clustal ⟨*Q*⟩ metric. The dominant main effect of the Urzyme accounts for >60% of the variance of the dependent variable, with the balance coming from the opposing effects of the B and C subsets (i.e., their β coefficients are of opposite sign). Regression models of the three independent metrics derived from BEAST 2 tree constructions all depend heavily on significant two-way interactions between segments of the different MSAs. Functional properties of the Class I aaRS modules are experimentally sufficiently well-characterized to sustain functional interpretations of the interaction terms for site-to-site metrics (Figs 2, 4) and an evolutionary interpretation of the row-by-row Support metric (Fig 5).

#### The relationship between Shape and segment B changes sign, depending on whether CP1 is present in the MSA

CP1 forms an annulus around one end of the two halves of the Urzyme (Fig 6A). Molecular dynamic simulations of the TrpRS Urzyme [17] show that in these two segments, which together form the amino acid binding site in Class I aaRS, exhibit extensive relative motion. Further, the distance between the two parts of the amino acid binding site decreases in the TrpRS catalytic conformational transition and that motion is coupled to the relative motion of CP1 and the anticodon-binding domain. Moreover, steady state kinetic measurements of amino acid specificity confirm that the relative domain motion is coupled to the relative specificity for Tryptophan vs Tyrosine [24].

**Fig 6.**
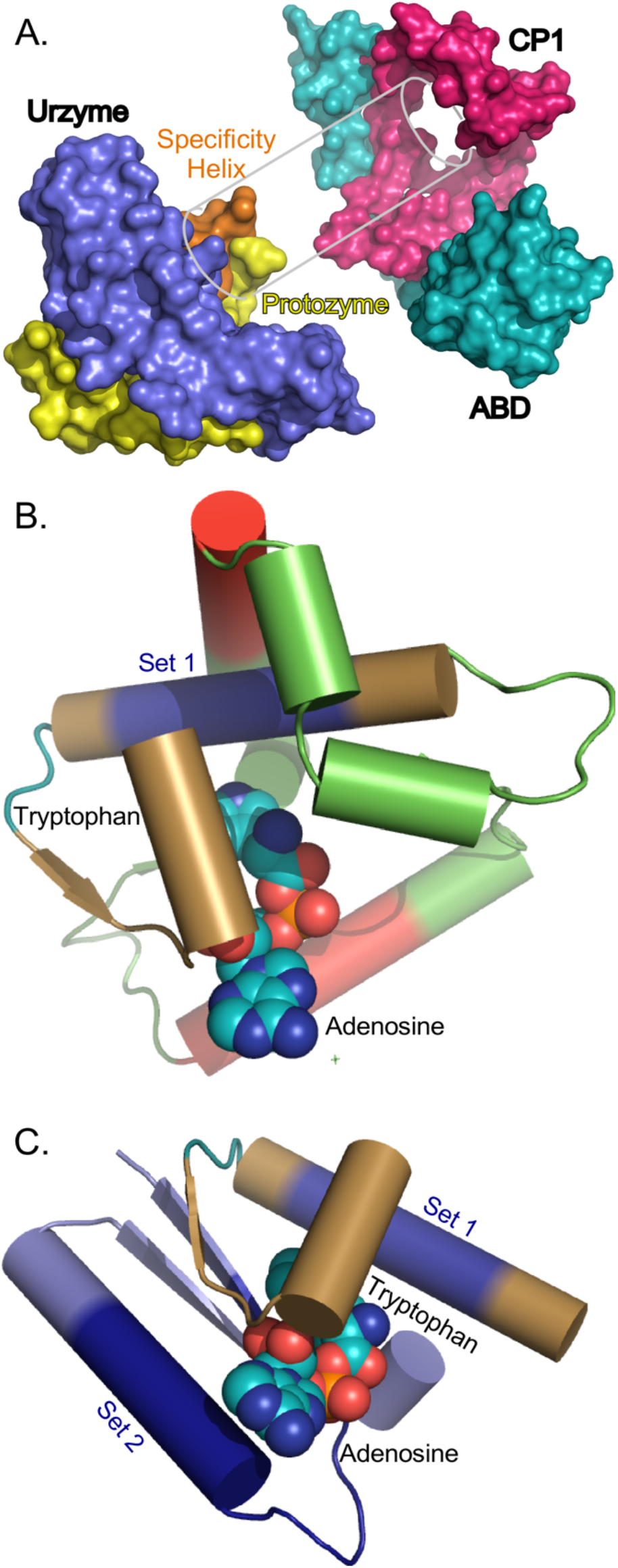
A. Structural relationships between the TrpRS Urzyme, CP1, and the anticodon-binding domain (ABD). Coloring in this Fig differs from that in Figs 1 and **B**. The CP1 motif forms an annulus that constrains motions of the specificity determining helix (sand) and the Protozyme (yellow), constraining the effective size of the amino acid binding pocket when the ABD (teal) changes its orientation. Experimental evidence described in the text confirms that these constraints enable full-length TrpRS to reject tyrosine in the transition state complex for tryptophan activation. **B**. Structural cartoon with details of the interaction illustrated in **A**. Coloring in this Fig is the same as that in Fig 1. The specificity-determining helix is the site of the Set 1 segment, which is colored in Blue. The TrpRS CP1 module is green and the scaffold segments are red. Note the close contact between CP1—especially the red scaffold segment—and Set 1. **C**. Interactions between the TrpRS protozyme—locus of the ATP binding site, and segment B, locus of the amino acid binding site. Subsets 2 and 1, which are contained entirely within the respective segments, are highlighted in dark blue.

Thus, the interaction between CP1 and the amino acid binding site of the Urzyme constrains the amino acid binding site dynamically. Structural relationships between CP1 and the amino acid biniding site in TrpRS are highlighted in Fig 6B. The experimental demonstration of the interaction between CP1 and the amino acid binding site validates the sign and strength of the B*CP1 contribution to Shape much as a pre-formed space in a partially assembled puzzle validates the Shape of the missing piece.

#### The relationship between Tree height and segment B changes sign, depending on whether the protozyme is present in the MSA

As noted in regard to Fig 2A,B, the Urzyme itself has a dominant impact increasing the Tree height, and this effect is reduced substantially by the protozyme but increased by the protozyme*Segment B interaction. Yet, sequences in Set 1 from within Segment B have the highest Tree height. Another reflection of the same phenomenon is that the protozyme and Urzyme appear in very much the same place on the regression plots in Figs 2 and 5, yet are quite well separated in Fig 4, where the protozyme is midway between the Urzyme and CP1.

The strong interaction in the Tree height regression model is between the protozyme—locus of ATP bnding—and segment B—locus of amino acid binding (Fig 6B, C). It is reasonable that the interaction term arises because of the contravariant effects of the protozyme, which has retained significant sequences over the evolution of the synthetases because it has the ATP binding site and Segment B, which has the amino acid binding site, and would be expected to exhibit the most significant evolutionary sequence variation between families specific for different amino acids.

#### The relationship between Support and CP1 changes sign, depending on whether the protozyme is present in the MSA

Residues from the protozyme contribute even more decisively to the average support than those derived from elsewhere in the Urzyme. CP1 has the most significant impact on the regression model for the Support metric. Its coefficient is -0.33, more than three times that of the next most important predictor. This effect can be seen in the regression plot in Fig 5, in which the CP1 MSA is more widely separated from the other MSAs than in the models for any other metric. However, the presence of the protozyme sequences in the Full MSA is sufficient to change the impact of CP1 from negative to positive, giving the β-coefficient of +0.1 for the protozyme*CP1 interaction term. This would appear to reinforce the conclusion, based also on the widespread occurrence of the protozyme packing motif [22] and its functional activation of ATP [21], that the protozyme was the original root of the entire superfamily and is older than the Urzyme itself, as previously proposed [46].

## Discussion

Our long-range goal is to understand how evolution of aaRS•tRNA cognate pairs effected the stepwise increases in dimensionality that resulted in the progression of entries into the universal genetic coding table. To that end we assembled three novel phylogenetic metrics here, to show that they identify high-resolution mosaicity in carefully curated, structure-based MSAs of secondary structures shared by all members of the Class I aaRS superfamily. All metrics support prior arguments that the Class I aaRS evolved by a succession of intermediates— protozyme=>Urzyme=>Catalytic domain—with increasingly sophisticated catalytic [14, 18, 19, 21, 31] and discriminatory [46-48] capabilities.

### Evolutionary refinement of protein catalysis was predicated on expanding the genetic code

The Tree height, Shape, and and Support metrics identify differences in the genetic origins of successively acquired contributors to aaRS function: protozyme=>Urzyme=>catalytic domain with CP1. They correlate inversely with experimentally measured catalytic proficiencies for the catalysts from which the corresponding sequences are derived (Fig 7A). Those proficiencies themselves correlate quantitatively with parallel increases in catalytic proficiency with the concomitant additions of mass in the Class I and II aaRS evolutionary intermediates they entail [See Fig. 6 in 14]. Both phylogenetic and functional properties of these intermediates thus appear to probe far deeper into evolutionary time than previous phylogenetic analyses of Class I aaRS [3, 7, 41-43].

**Fig 7.**
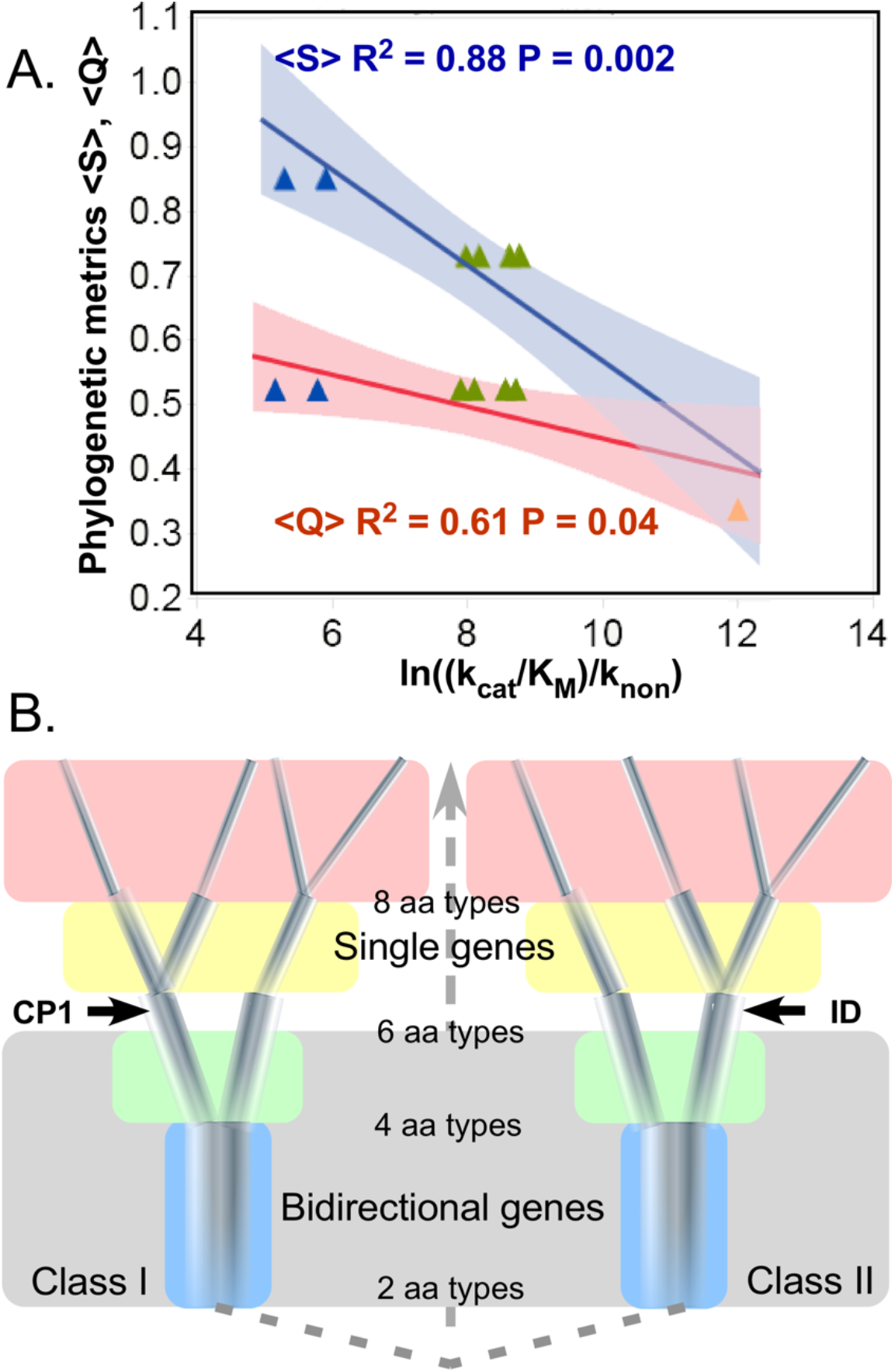
Assignment catalysis and code evolution. **A**. Correlations between rate accelerations and phylogenetic metrics, ⟨*Q*⟩ and ⟨*S*⟩. Parameters for regression against experimental catalytic proficiencies for corresponding putative evolutionary intermediate constructs are both significant. Blue, green, and amber triangles represent protozymes, Urzymes, and catalytic domains. **B**. Timeline for growth of the genetic coding alphabet from a two-letter code. Introducing new aaRS into the context of the ancestral bidirectional gene (grey background and dashed connecting lines) simultaneously enhanced specificity and created fundamental changes in selection pressure. Different colored backgrounds signify altered selection pressures that apply to all extant aaRS at a given stage of the coding alphabet. Increasing cardinality of the alphabet induces (i) sequence space inflation as a greater number of distinguishable sequences can be specified; and (ii) restriction in the diameters of the quasispecies as they approach fully coded sequences in the final 20-letter alphabet. Bold face landmarks (CP1 and ID) denote qualitative changes in aaRS architecture shown experimentally to enrich specificity as discussed in the text. Note that this timeline like that in Fig 5 represents evolutionary events before the LUCA.

The precision with which protein active sites distinguish substrates from one another and transition states from substrates was the result of the evolutionary process we wish to characterize. Expansion of genetic coding itself depended critically on developing a system for the placement of a precisely defined amino acid sidechain at a particular point on a peptide backbone. Phylogenetic analysis of segmental Class I aaRS MSAs represents a uniquely promising opportunity to test the hypothesis that contemporary enzymes are mosaic structures rooted in simpler catalytic polypeptides and assembled from detectably different genetic ancestors.

### High-resolution structural modularity implies functionally significant discontinuities in the evolution of genetic coding

If CP1 insertions were indeed assimilated from one or more similar genetic sources after Class I aaRS Urzymes had evolved significantly from modest ancestral protozymes, it could have at least three noteworthy biological implications:

i. Class I protozymes, whose high catalytic activity mobilizes ATP for biosynthesis, an activity found in many proteins—may be the root of that major portion of the proteome built from β-α-β crossover connections—the Rossmannoid protein superfamily [22] and potentially β-barrel proteins [49, 50], which are central to intermediary metabolism and nucleotide biosynthesis [51].
ii. AARS protozyme and Urzyme populations would have functioned first as quasispecies in translation, limiting the sophistication of the early proteome.
iii. CP1 assimilations would have transformed selection pressures for subsequent aaRS evolution by facilitating enhanced fidelity.

#### Primordial aaRS quasispecies covered progressively smaller regions of sequence space, closer to the “root”

Early aaRS evolution cannot have been an ordinary mutation/selection process. All solutions to the problem normally framed as a “chicken and egg problem” [52] imply historical context and a continuity principle, which must be defined by the genotype-phenotype mapping [53]: at any stage during the emergence of coding. Some prior system must have been interpreted extant aaRS protogenes. A key aspect of such prior systems is that as the alphabet size, diversity, and consequent fidelity increased, they would, *ipso facto*, have created more narrowly targeted selection pressures, strongly coupling mutation and selection in early stages of genetic coding (Fig 7B). The subsequent coevolution of CP1 and Urzyme sequences would necessarily have preserved Urzyme functionality, while the new CP1 could adapt flexibly to its developing role of enhancing specificity as described in the next section.

Different background colors in Fig 7B denote how branching of the tree to allow introduction of the *n*^th^ amino acid into the alphabet enforces a highly cooperative re-optimizing of the *n*–1 aaRS types already present. As each new, refined amino acid type emerged, all extant gene sequences adapted to opportunities introduced by progressively finer discrimination between amino acid side chain physical chemistry. In turn, adaptation to a more diverse alphabet sharpens the new aaRS fidelities (i.e., the contraction of branch thicknesses in Fig 7B), implicating a bootstrapping feedback and enhancing the cooperativity of the transition to higher-dimension coding alphabets [46]. That cooperativity creates a Lamarckian correlation between selection pressure and its outcome—the result of any mutation being nearly synonymous with the selection pressure it faced, especially as the code differentiated. Survival would have depended on the relationship between the shapes of fitness landscapes and error rates of catalysis by the extant quasispecies [54]. The earliest alphabets, including at least the first two amino acid types, were also tightly constrained by bidirectional coding (gray shaded background, Fig 7B).

We have argued [46, 48] that a single functional island in sequence space (i.e. quasispecies) would invariably have been a strong attractor, irrespective of the detailed Shape of the fitness landscape that stabilized it, because all mutations that moved the system slightly off its optimum phenotype would be subject to strongly restorative selection pressures. However, if ancestral protozymes from a bidirectional gene had broad, relatively flat (and necessarily co-dependent) fitness landscapes, matched to correspondingly high error rates, that could have favored bifurcated quasispecies that enhanced genetic coding by recruiting new amino acid types and simultaneously increasing the precision with which child specificities could be encoded.

#### Insertion of CP1 likely enabled saltatory improvements in fidelity

Structural and biochemical data suggest that the CP1 insertions created stepwise enhancements in the evolution of genetic coding by enabling conformationally-driven mechanisms to enhance specificity. The shortest CP1 insertions have ∼75 residues in TrpRS and TyrRS that recur essentially intact in the longer CP1 insertions of the remaining eight Class I aaRS [30, 36]. It seems likely that the initial insertion needed to be that long. CP1 forms an annulus around the Urzyme (Fig 6), constraining relative movements of the protozyme and specificity helix that form the amino acid binding site [17]. For that reason, the original hypothesis as to its origin was near-simultaneous insertion of a mobile genetic element into all extant Class I Urzymes [30], in which case its root sequence would have been more recent than that of the Urzymes, yet earlier than contemporary full-length aaRS enzymes.

Amino acid specificities [18, 19] suggest that although capable of 10^9^-fold rate accelerations, Class I and II Urzymes select an amino acid from the correct Class only 80% of the time. However, Wills & Carter [20] note that their within-Class specificity is consistent with each Class distinguishing two kinds of amino acids, to operate a four-letter coding alphabet. These modest fidelities suggest a fundamental limit to the precision of which bidirectional coding was ultimately capable.

The most evident contribution of CP1 to fidelity is that the editing domains present in the larger subclass IA aaRS for aliphatic amino acids Ile, Val, and Leu are elaborations of the CP1 motif present in the simplest subclass IC aaRS for Tyr and Trp. It seems likely, however, that CP1 functioned earlier to enhance fidelity by dynamically constraining the size and Shape of the amino acid binding pocket. Several groups found that comprehensive mutation of side chains in the immediate vicinity of the amino acid substrate, all within the Urzyme architecture, would not change specificities of subclass IB GlnRS to Glu [55, 56] or subclass IC TrpRS to Tyr [57]. Changing GlnRS specificity to Glutamate (Bullock, et al. [56]) required wholesale mutations in the second layer surrounding the amino acid binding site outside the Urzyme, but within in the GlnRS CP1 domain.

Similarly, a modular thermodynamic cycle comparing specificities of full-length TrpRS, Urzyme and Urzyme plus either CP1 or the anticodon binding domain (ABD) showed rejection of Tyrosine by *G. stearothermophilus* TrpRS requires cooperation between CP1 and the ABD [24]. CP1 must coordinate its movements with those of the ABD to perform the allosteric communication necessary to enhance side chain selectivity beyond the modest capabilities of the Urzyme [25, 27]. In both cases, fine tuning specific recognition of amino acid substrates required insertion of CP1 and the ABD and, subsequently, coupling between them.

Inserting the CP1 motif would necessarily have disrupted bidirectional coding (Fig 1B), thus dividing aaRS evolutionary histories decisively into distinct stages (Fig 7, between green and yellow backgrounds). Selective advantages of CP1 insertion thus appear to have been to (i) end constraints imposed by bidirectional coding and (ii) transcend the fundamental limitation on specificity posed by the Urzyme architecture. Either or both would have allowed substantial increases in fidelity to develop. CP1 therefore likely dramatically transformed the Class I aaRS fitness landscape, and was likely necessary to expand the coding alphabet.

#### A revised branching order suggests that Class I aaRS protozymes and Urzymes root the Rossmannoid superfamily

Putting Class I aaRS protozymes at the root of that superfamily reconfigures many branching orders within the proteome to reflect that aaRS Urzymes were not a late-developing branch in the Rossmannoid superfamily radiation, but instead were ancestral to it (Fig 8). Descent of Class I and II aaRS from a single bidirectional ancestral gene [10, 14, 18, 19, 21, 32, 33] would underscore the likelihood that the aaRS of both families diverged, rather than converging to similar functions from different sources. Thus, the genetic coding table itself likely evolved by bifurcating pre-existing aaRS genes into specialized enzymes whose more discriminating specificities for tRNA and amino acid substrates enabled daughter synthetases to differentiate groups amino acids that previously had functioned as synonymous [46, 48, 58, 59]. It would be surprising if other branches of the proteome had not diverged from the ancestral aaRS, as suggested in Fig 6.

**Fig 8.**
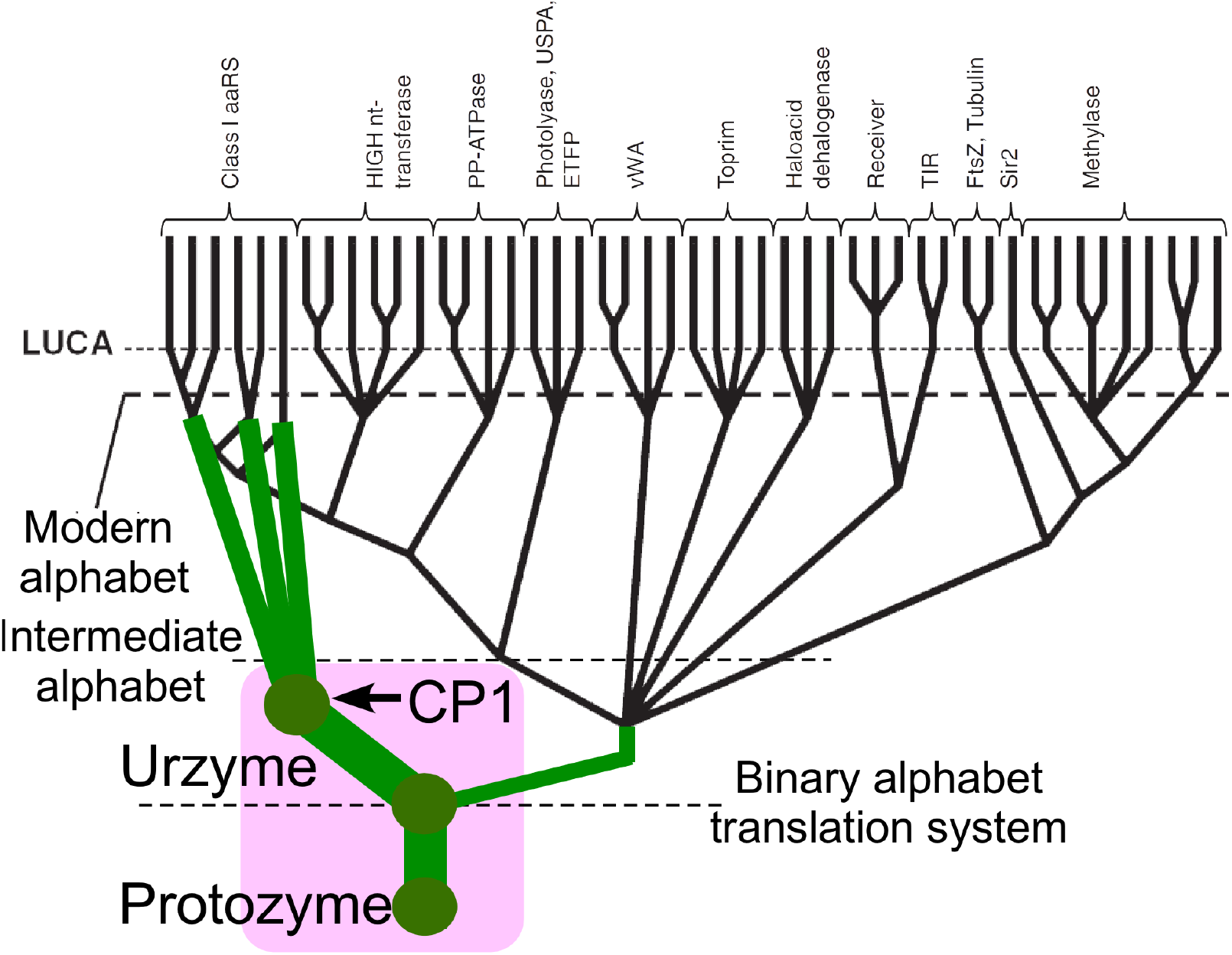
Modified Class I aaRS phylogeny consistent with bidirectional coding ancestry in a peptide/RNA world. Adapted with permission from Fig 12-2A of [62]. Green circles represent key stages in the emergence of genetic coding, beginning with ATP-dependent amino acid activation by protozymes coded by a bidirectional gene [21]. Magenta background indicates the progression illustrated in detail in Fig7B.

### Phylogenetic trees have meaningful fine structure

The coherence of the phylogenetic metrics described in previous sections makes a strong case for integrating complementary experimental data into a coherent application of Bayesian phylogenetic analysis. Their functional significance emerges only in the context of partitioning the overall MSA according to structural and functional criteria established with other disciplines and in the presence of suitable controls for the effect of sequence length (subsets 0-6 from the Urzyme MSA; Fig 1C, Table 1). The evidence that emerges for fine structure within protein domains is without precedent.

In turn, the phylogenetic analysis provides novel validation of the experimental work that guided decisions on partitioning the MSA. Identification of how the Tree height and Shape metrics depend on intermodular two-way interactions validates the considerable experimental work demonstrating how intermodular coupling contributes to catalysis and specificity. Further, because these interactions arise from the coherence of the entire Class I superfamily, they imply that much of that experimental evidence applies, perforce, to similar interactions between ATP and amino acid binding sites and between amino acid binding sites and CP1 in all or most Class I aaRS.

These observations suggest that this work points the way to more substantial applications of software like BEAST2 [60] to a broader range of evolutionary questions.

### Genetic coding and the proteome probably emerged from an RNA•polypeptide partnership

Previously accepted phylogenies of the Class I aaRS root well before the Last Universal Common Ancestor (LUCA) [3, 7, 41-43], but appeared also to have radiated after earlier bifurcations in the immense meta-family [3] containing folds based on the Rossmann dinucleotide-binding fold [61]. This paradoxically late radiation of protein assignment catalysts is the foremost phylogenetic evidence favoring the RNA World hypothesis. Genetic coding by protein aaRS was therefore thought, necessarily. to have replaced a prior implementation based on ribozymal assignment catalysts [62, 63].

Evidence described here that CP1 is a much more recent acquisition by more ancestral aaRS Urzymes supports a plausible alternative branching order (Fig 8). Attributing the apparently late adaptive radiation of Class I aaRS to CP1 achieves consistency with an origin of genetic coding from a bidirectional gene administering a binary coding alphabet in a peptide/RNA world. Resolving the paradox in this way complements—without necessarily contradicting—the phylogenetic analyses of Koonin [62, 63].

## Methods

Amino acid chemistry underlies the metric form of amino acid substitution matrices (aaSM) generally required for the logically circular process of creating an amino acid sequence alignment by optimizing the constraints of some aaSM and then building a phylogenetic tree from the alignment according to those constraints [8].

We superposed the available bacterial Class I aaRS crystal structures to base the multiple sequence alignment (MSA) exclusively on precise three-dimensional structural homology, freeing the MSA from dependence on empirical substitution matrices and estimates of relative rates of mutation [64]. In the first place, this was done within each aaRS type. The identification of sequence positions displaying very high conservation of both structure and amino acid occupancy provided unambiguous anchors for the production of much more extensive sequence alignments. Popinga [65] performed this task through extensive use of HHpred [66] and MODELLER [67] to thread amino acid sequences for each aaRS type using experimentally determined 3D structures as templates. This procedure produced, for each aaRS type, a greatly expanded MSA in which various well-conserved secondary structural regions were near-perfectly structure- and sequence-aligned between species known to have diverged not long after the LUCA.

In the final stage of alignment, the structures of the individual aaRS types were brought together to identify regions of universal structural homology, the Class I aaRS “scaffold”. The product was an MSA across all Class I aaRS types in which each sequence position could be assumed to have arisen, as far as is reasonably possible, from the same LUCA-ancestral codon. While the validity of this assumption is by its very nature untestable, the close structural homology of Class I aaRS of all types and the rigour of our procedure gives us confidence and justification as good as that underlying any cross-species phylogenetic enterprise. However, we took further steps to restrict and constrain the data upon which we later built putative phylogenies. First, individual scaffold elements are more extensive in some extant aaRS types than others and it is impossible to identify any proper universal homology among the loops, turns and structurally disordered regions that join them. All of these were excluded from our grand Class I “scaffold” MSA. Second, we included only eubacterial aaRS. It has been suggested previously [10] that bacterial aaRS sequences are closer to the ancestral root than those from other domains. While analysis of loop structures demonstrates that this is contestable in respect of the ages of biology’s three main evolutionary domains (ref Berezovsky? Lupas? Trifonov?) it is generally clear that cytoplasmic proteins of bacteria have been subject to many fewer complex selection pressures than their archeal and eukaryotic counterparts. The complex functional and regulatory roles played by some aaRS proteins and their involvement in numerous genetic syndromes attests to this (ref).

Thus, the final MSA contained roughly bacterial 20 sequences for each of 10 Class I aaRS, circumventing as many problems as possible in aligning sequences that diverged to produce different substrate specificities in the pre-LUCA æon. The use of scaffold sequences for phylogenetic analysis gave the best guarantee that results would reflect relatively neutral evolution within the context of asymmetric, specialised selection pressures producing variant substrate specificities by fine-tuning the size, Shape and chemistry of more intricately constructed amino acid side-chain structures and pockets.

The MSA for the Class I scaffold was output using VMD [68], and is provided in the supplement (Supplement_MSA_files.txt) together with all subsets used in this work. The scaffold fasta file was then partitioned along lines of the experimental deconstruction of the Class I aaRS superfamily [14] into the disjointed segments of the Urzyme, separated by structurally conserved segments from CP1. Finally, because the Urzyme (83) and CP1 (20) partitions of the scaffold MSAs have different sequence lengths, seven subsets comprising 20 sequence positions distributed throughout the Urzyme were selected arbitrarily by a balanced, randomized procedure [69] to test the effect of sequence length on our phylogenetic signatures.

We compiled a three-dimensional comparison of complementary metrics of the overall scaffold MSA, the full Class I Urzyme, the protozyme, and CP1 subsets. BEAST 2 accumulates two metrics from the ensemble of trees generated by Markov Chain Monte Carlo simulation. The Tree height reflects the mean overall mutation rates, and site-specific mutation rates are fitted to a gamma distribution [70] by adjusting a Shape parameter, α = 1/CV^2^, where CV is the coefficient of variation or the ratio of the standard deviation to the mean and thus a measure of Tree height variance.

For the third dimension, molecular kinships between rows of the MSA within each partition were compared by clustering the sequences into clades according to their evolutionary origin. Phylogenetic trees were computed using BEAST2 [60], allowing the use of multiple amino acid substitution matrices (primarily WAG and LG). Trees were visualized with DensiTree [71] and FigTree [72]. For the full Class I scaffold and each partition of interest, DensiTree was used to extract ∼10,000 trees generated by BEAST2 in building trees. From these we calculated values of a parameter representing the support for clades that each corresponded exclusively to an individual aaRS type, *i*. This support, *S*_*i*_, is the percentage of all trees for which the aaRS type in question appears to be monophyletic, meaning that the leaves of that aaRS type descend from the same most recent branch point (common ancestor) with no descendants arising from a different aaRS type. The mean phylogenetic support, ⟨*S*⟩, is a metric derived from a row-by-row calculation over all 10,000 phylogenetic trees generated using the sequence data for a chosen partition of the scaffold MSA.

The conservation quality, ⟨*Q*⟩_j_, defined by Clustal [37, 38] was computed down each column, *j*, of the grand MSA, which included all aaRS types. While the definition of ⟨*Q*⟩ seems convoluted, it has been constructed as a metric whose value reflects the degree of amino acid diversity generated by typical evolutionary amino acid substitution processes, reflected in the substitution matrix that it uses as a reference. Different matrices do not give widely divergent ⟨*Q*⟩ parameters. We used the ClustalW default matrix (PAM 250; [73]) and calculated the average, ⟨*Q*⟩, over all positions within a partition of the MSA a parametric representation of the partition-wide variation in amino acid occupancy calculated column-by-column over individual sequence positions.

Columns for various predictors were appended to the table of ⟨*Q*⟩ and ⟨*S*⟩ values to form the design matrix (Table 2) used for multiple regression analysis with the JMP software [74]. Multiple Regression was used to assess the statistical strength of contributions of the predictors (i.e., the independent variables, A, B, C, CP1 defined in Fig 1 and the number of amino acids) to ⟨*Q*⟩ and ⟨*S*⟩ values, using stepwise searches to identify the best set of predictors followed by least squares estimation of the corresponding coefficients and their Student t–test P values.

## Abbreviations

aaRS: aminoacyl-tRNA synthetase;
CP1: Connecting peptide 1;
MSA: multiple sequence alignment;
MCMC: Markov chain monte carlo

## Acknowledgments

This work was supported by NIGMS (Grant number R01-78227 to C.W.C., Jr). We are grateful to Corbin Jones, Greg Fournier, Günter Wachtershäuser, and Bill Martin for helpful feedback in preparing the manuscript and Loren Williams for suggesting that ribosomal protein sequences may exhibit a similar genetic mosaicity.

